# OsCYP71D8L as a key regulator involved in growth and stress response by mediating gibberellins and cytokinins homeostasis in rice

**DOI:** 10.1101/538785

**Authors:** Jiahao Zhou, Zeyu Li, Guiqing Xiao, Rongfeng Huang, Haiwen Zhang

**Author notes:** Correspondence authors: E-mail, Haiwen Zhang,. Rongfeng Huang,. These authors contributed equally to the article (Jiahao Zhou, Zeyu Li).

## Abstract

Phytohormones are pivotal in the regulation of plant growth and development, and acclimation to adverse environments. Multiple cytochrome P450 monooxygenases (CYP450s) are involved in the biosynthesis and catabolism of phytohormones. Here, we reported that a CYP450 member of CYP71 clan, OsCYP71D8L, participated in the control of multiple agronomic traits and abiotic stress responses by affecting gibberellin (GA) and cytokinin (CK) homeostasis in rice. We found that its gain-of-function mutant (*cyp71d8l*) and transgenic plants overexpressing *OsCYP71D8L* (*OsCYP71D8L*-OE) displayed similar phenotypes such as dwarfed plant, reduced panicle length and grain number per panicle. In *OsCYP71D8L*-OE seedlings, endogenous GAs and CKs was notably decreased and increased as compared with wild type (WT), respectively. Correspondingly, the dwarfed plant and less developed root of *cyp71d8l* and *OsCYP71D8L*-OE seedlings could be rescued by exogenous GA3, but more exacerbated by exogenous 6-BA. Importantly, *cyp71d8l* and *OsCYP71D8L*-OE seedlings maintained high chlorophyll contents and low reactive oxygen species level, and showed significantly enhanced tolerances to drought and high salt compared with WT. Thus, our results suggest that OsCYP71D8L plays roles in regulating rice growth and stress responses by coordinating GAs and CKs homeostasis, and it is useful to engineer stress-tolerant rice.

## Introduction

Rice (Oryza sativa L.) is an important food crop for over half of the human population, but various environmental stresses including high salinity, drought, and extreme temperature, seriously affect the developmental processes and productivity of rice (*Oryza sativa*). Thus, there is a strong demand for developing abiotic stress-tolerant rice to ensure food supplies for the increasing population (Mizoi et al. 2013). Plant height is a key agronomic trait determining plant architecture, yield, and stress response in rice, which is controlled by phytohormones and their intergraded signal networks (Liu et al., 2018). Many genes involved in gibberellin (GA) biosynthesis and signal including *SD1, GDD1, SLR1, EUI, GID1, SD1* and *D1* were reported to play crucial roles in plant height, architecture, organ development, yield and stress response in rice (Liu et al. 2018; Kwon, C.T. and Paek, N.C. 2016; Colebrook et al. 2014; Li et al., 2011). For examples, manipulating GA levels by ectopic expressing modified *GA2ox6* led to the increased grain yield, dwarfed plant, more tillers, expanded root system, and the elevated drought and disease tolerance (Lo et al., 2017). Ectopic expression of *PtCYP714A3* resulted in the lower GA level associated with semi-dwarfed plant, reduced seed size, and enhanced salt tolerance (Wang et al., 2016). Moreover, GA-insensitive dwarf mutant *gid1* showed increased tolerance to various abiotic stresses including cold, drought and submergence by maintaining high levels of chlorophyll and carbohydrates, and low reactive oxygen species (ROS) accumulation under stress conditions (Tanaka et al., 2006).

Cytochrome P450s are the largest protein family members catalyzing many reactions involved in the biosynthesis and catabolism of phytohormones, pigments, antioxidants and defense compounds (Renault et al., 2014). Recently, multiple P450s have been reported to participate in the regulation of developmental processes and stress response by modulating GA homoeostasis in different plants. In *Arabidopsis*, two members of CYP88A, KAO1 and KAO2, overlappingly regulated GA biosynthesis throughout plant development (Regnault et al., 2014). Two EUI homologs, CYP714A1 and CYP714A2 were involved in regulation of plant growth and development by deactivating of bioactive GAs (Nomura et al., 2013; Zhang et al., 2011). In rice, two paralogs of CYP701, OsKOL4 and OsKOL2, acted as ent-kaurene oxidase involved in gibberellin biosynthesis (Itoh et al., 2004; Wang et al., 2012). EUI1/CYP714D1, CYP714B1 and CYP714B2 deactivated bioactive GAs and resulted in the elongated uppermost internodes (Magome et al., 2013; Zhang et al., 2008; Zhu et al., 2006). Moreover, overexpression of *DSS1* led to reduced GA and increased ABA associated with the dwarf phenotype (Tamiru et al., 2015).

The rice genome contains at least 356 P450 genes, which are grouped into the 10 plant P450 clans. The CYP71 clan is the largest set of P450s and represents more than half of all CYPs in higher plants (Nelson and Werck-Reichhart, 2011; Nelson et al., 2004). In plants, CYP71BL1, CYP71DD6 and CYP71AM1 are involved in the synthesis of eupatolide, inunolide, and sorgoleone, respectively (Frey et al., 2018; Pan et al., 2018). We previous showed that *CYP71D8* was differentially expressed in two rice varieties under drought stress (Zhang et al., 2012), but the functions of CYP71 members remain unknown in rice. Here, we reported that a CYP450 member of CYP71 clan, OsCYP71D8L, participated in the control of multiple agronomic traits and abiotic stress responses by affecting GAs and cytokinin (CK) homeostasis in rice. Our results suggest that OsCYP71D8L might act as a key regulator in balancing plant growth and development process, and stress responses by affecting hormone homeostasis in rice

## Materials and methods

### Plant materials and phenotype analyses

Japonica rice (*Oryza sativa* L.) Koshihikari (KOS), *OsCYP71D8L*-OE lines, Hwayeong (HY), and a T-DNA insertion mutant of *OsCYP71D8L* (*cyp71d8l*) ordered from the Crop Biotech Institute at Kyung Hee University were used in this study.

All plants grown in pots with soil were used to determine the stress tolerance. For dehydration treatment, 3-week-old plants were subjected to drought stress for 10 days by withholding water and then re-watering for 7 days. For treatment with salt stress, 3-week-old plants were watered with 150 mM NaCl for 7 days and then recovered normal growth for another 7 days. When the seedlings began to show stress symptoms (i.e. leaf rolling), phenotypes of plants from two stress treatments were observed and analyzed.

All rice materials were planted in a field during the natural growing season for the analyses of agronomic traits including plant height, internode length, internode number, panicle length, and grain number per panicle.

### Phylogenetic analysis

The putative homologs of OsCYP71D8L from different monocot plants were identified using the Basic Local Alignment Search Tool from the National Center for Biotechnology Information (http://blast.ncbi.nlm.nih.gov/Blast.cgi). A phylogenetic tree of OsCYP71D8L homologs was constructed using the fast-minimum evolution algorithm (https://www.ncbi.nlm.nih.gov/blast/treeview/).

### Cell length observation

The uppermost internodes of KOS and *OsCYP71D8L*-OE at the maturation stage were fixed in 2.5 % v/v glutaraldehyde and then dehydrated through a series of graded ethanol concentrations. The samples were embedded in resin and sectioned. Sections were stained with toluidine blue, and then microscopically photographed.

### Determination of chlorophyll, soluble sugar and H_2_O_2_ contents under stress

Three-week-old seedlings of KOS, *OsCYP71D8L*-OE lines, HY and *cyp71d8l* were subjected to drought for 5 days or 1% NaCl treatment for 3 days, respectively. Then, the contents of chlorophyll, soluble sugar and H_2_O_2_ in the leaves of all materials were measured as reported previously (Urano et al., 2014, Song et al., 2011), respectively.

### Determination of GAs and CKs

Endogenous GAs and CKs were measured using an LC-ESI-MS/MS system (Chen et al. 2013). About 100 mg fresh leaves of 4-week-old KOS and *OsCYP71D8L*-OE seedlings were harvested and ground into powder in liquid nitrogen. Then, all materials were extracted with 1 mL methanol containing 20% water at 4°C for 12 h. After centrifuged at 12,000 g under 4 °C for 15 min, the supernatant was collected and evaporated to dryness under nitrogen gas stream. After reconstituted in 100 mL acetonitrile containing 5% water, the contents of GAs and CKs in all sample extracts were analyzed by LC-ESI-MS/MS system.

### Exogenous GA and 6-BA treatment

For GA and 6-BA treatments, more 20 germinated seeds of KOS, *OsCYP71D8L*-OE lines, HY and *cyp71d8*l were hydroponically cultured with 0.1 and 1 μM GA3, or 0.001, 0.1 and 1 μM 6-BA, respectively. After 6 days, the plant height and root growth were observed.

## Results

### OsCYP71D8L is a member of CYP71 clan

Intriguingly, the rice genome contains multiple paralogs of CYP450 as an eight-gene tandem array on chromosome 2 (Fig. 1A). Among them, LOC4328530 (OsCYP71D7), LOC4328531 (OsCYP71D8L) and LOC4328532 (OsCYP71D8) were belonged to CYP71 clan. Phylogenetic analysis showed that OsCYP71D8L was more closely clustered with three rice proteins (OsCYP71D8, LOC4328535 and LOC4328533) than the homologs from the other monocot plants (Fig. 1B). Moreover, five clustered proteins (OsCYP71D8L, OsCYP71D8, LOC4328533, LOC4328535 and OsCYP71D7) shared higher degree of identities. Particularly, OsCYP71D8L and OsCYP71D8 had 91.19% amino acid identity (Fig. S1). Of note, CYP71D8 was differentially expressed in two rice varieties under drought (Zhang et al., 2012), but the function of neither OsCYP71D8L nor OsCYP71D8 is known.

**Figure 1.**
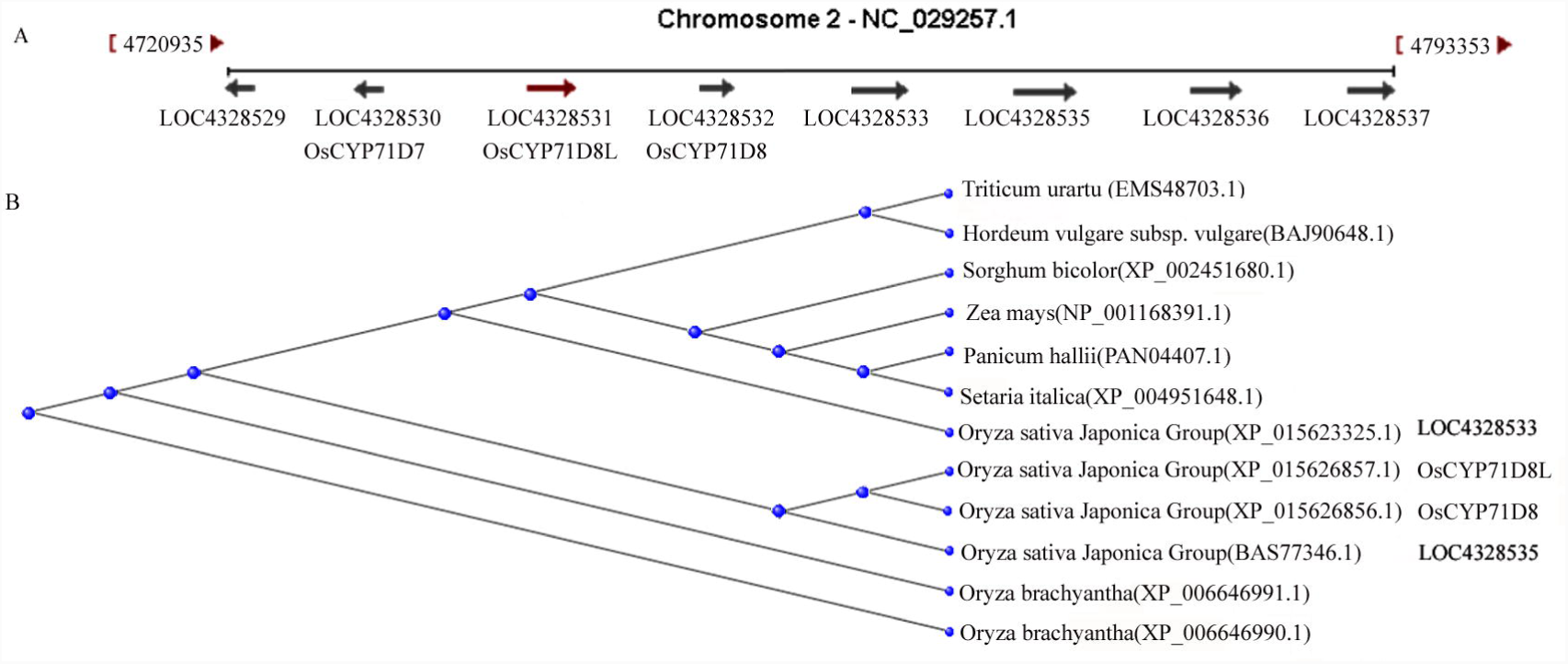
Tandem CYP450 paralogs and phylogenetic analysis of OsCYP71D8L. A, Tandem array of eight CYP450 genes on chromosome 2 (image obtained from NCBI database). B. Phylogenetic relationships of OsCYP71D8L with the homologs from different monocot plants, including OsCYP71D8L, CYP45071D8, LOC4328533 and LOC4328535 of *Oryza sativa Japonica*, NP_001168391.1 of Zea mays, XP_006646990.1 and XP_006646991.1 of *Oryza brachyantha*, BAJ90648.1of *Hordeum vulgare subsp. Vulgare*, XP_004951648.1 of *Setaria italica,* PAN04407.1 of *Panicum hallii*, EMS48703.1 of *Triticum Urartu*, and XP_002451680.1 of *Sorghum bicolor*. The fast-minimum evolution tree with the query ID of XP_015626857.1 was produced using BLAST pairwise alignments (https://www.ncbi.nlm.nih.gov/blast/treeview).

### *OsCYP71D8L* affects multiple agronomic traits in rice

For the possible functional characterization, we generated the transgenic plants overexpressing *OsCYP71D8L* (*OsCYP71D8L-OE*) driven by the C*aMV 35S* promoter. We found that *OsCYP71D8L-OE* lines displayed a severe dwarf phenotype throughout the life cycle (Fig.2A, Fig. S3A-C). The plant heights of *OsCYP71D8L-OE* lines were about 62-66% that of WT (KOS) at the mature stage (Fig.3A-B). Moreover, the panicle length and grain number per panicle of *OsCYP71D8L-OE* lines were decreased to about 90% and 66% of KOS (Fig.2B). We confirmed a homozygous T-DNA insertion mutant of *OsCYP71D8L* (*PFG_1C-03701.R*, named *cyp71d8l*) by PCR test. Sequencing analysis indicated that the T-DNA was inserted in the 2^nd^ exon of *OsCYP71D8L* and caused the truncated protein with 30 amino acids deleted in the C-terminus (Figure S2 A-C). *cyp71d8l* showed no notable difference with HY in vegetative growth stage (Fig.S3A-B). However, it began to exhibit dwarf phenotypes after flowering stage (Fig S3C), and showed reduced plant height (80%) (Fig.2A, 3A-B), panicle length (96%) and grain per panicle (78%) (Fig.2B) in comparison with HY at the mature stage, suggesting that *cyp71d8l* is a gain-of-function mutant with the phenotypes like that of *OsCYP71D8L-OE* lines. These results implied that OsCYP71D8L plays pleiotropic roles in regulating plant height and yield-relater traits.

**Figure 2.**
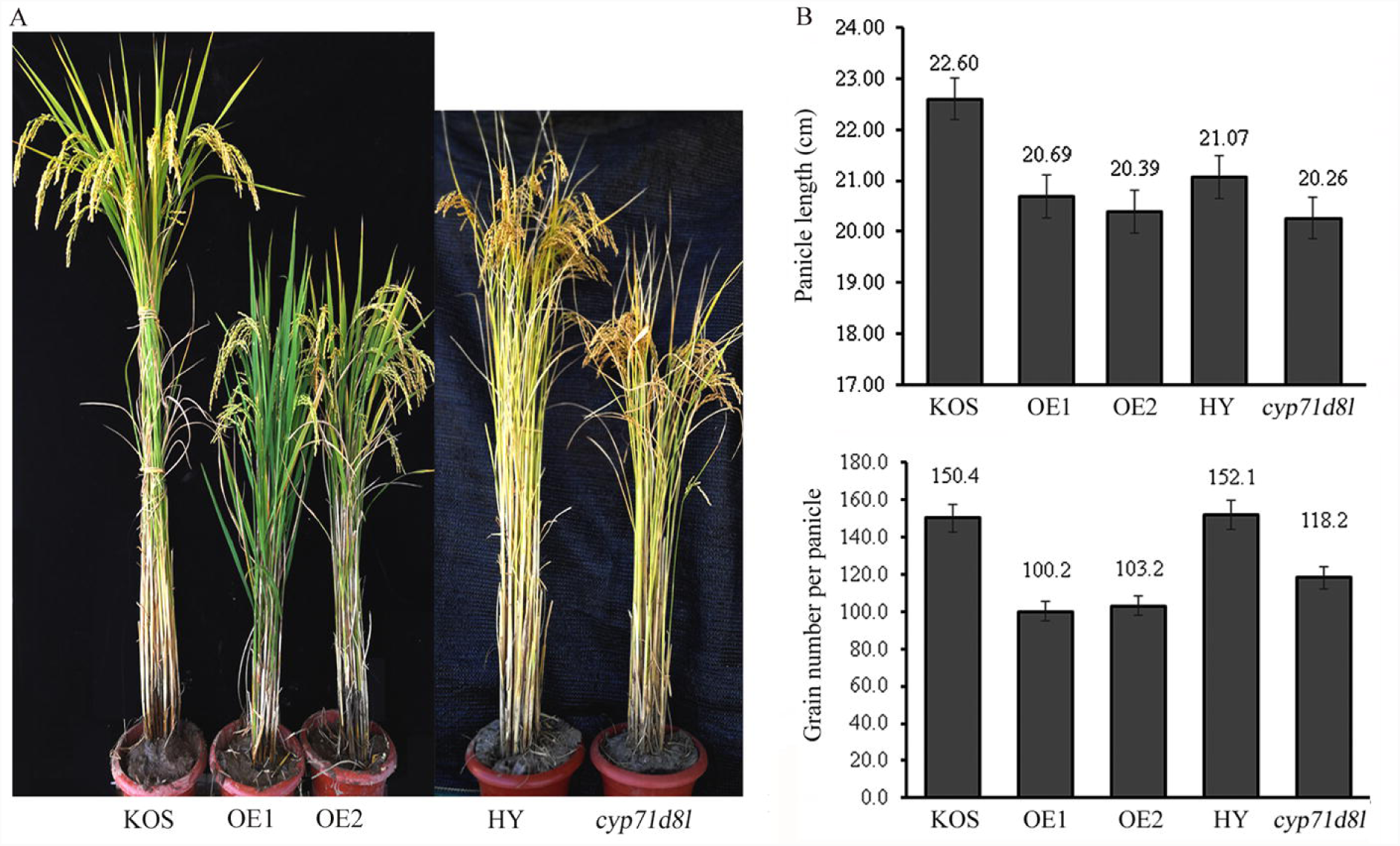
OsCYP71D8L affects plant height, the length of panicle and grain number per panicle. A, The phenotype of KOS, *OsCYP71D8L*-OE, HY and *cyp71d8l* at the at the mature stage. B, The panicle length and grain number per panicle of KOS, *OsCYP71D8L*-OE, HY and *cyp71d8l*. Data are average of at least 15 plants.

**Figure 3.**
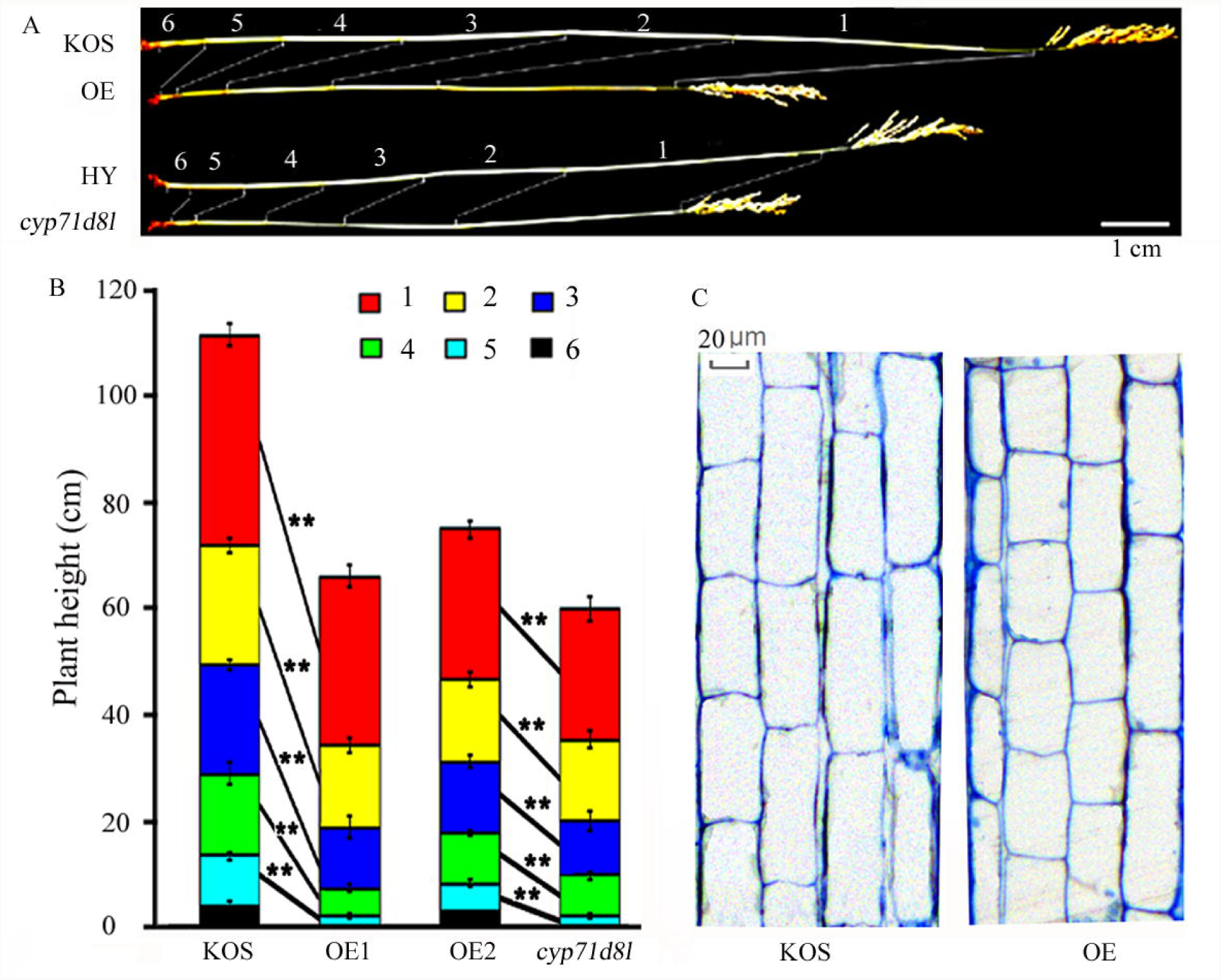
Reduced culm and internode length in *OsCYP71D8L-OE* and *cyp71d8l* plants at the mature stage. A, Representative culm lengths of KOS, *OsCYP71D8L*-OE, HY and *cyp71d8l* plants. Internodes are numbered from top to bottom as described. Bar=10 cm. B, Length of the first to sixth internodes in KOS, *OsCYP71D8L*-OE, HY and *cyp71d8l* plants. Internode positions are numbered as in (**A**). Mean and standard deviation for each internode are presented at least 15 plants. Asterisks indicate significant difference at P <0.01 compared with the wild type by Student’s t test. C, Parenchyma cell length of the uppermost internode in KOS and *OsCYP71D8L*-OE plants. Longitudinal tissue sections were stained with 0.3 % toluidine blue and observed under microscope. Bar=20 μm.

### OsCYP71D8L affects rice nodes number and internode length

In rice, the node number and internode length are the major determinants of plant height. We found that the dwarf phenotypes of *OsCYP71D8L-OE* and *cyp71d8l* were caused by reduced node numbers and internode length (Fig.3A), *OsCYP71D8L-OE* and *cyp71d8l* plants developed only 5 nodes, while WT plants developed 6 nodes. Moreover, the average internode length of *OsCYP71D8L*-OE and *cyp71d8l* was significantly shorter than that of KOS and HY(*P*<0.01), respectively (Fig. 3B). The decreased internode length could be caused by the cell proliferation or cell elongation. Our microscopy analysis revealed that the parenchyma cells in the uppermost internode of *OsCYP71D8L*-OE at the mature stage was clearly shorter that of KOS (Fig 3C), indicating that the reduced internode number and internode length account for the dwarf phenotypes of *OsCYP71D8L-*OE and *cyp71d8l*.

### OsCYP71D8L affects gibberellins homeostasis

Since *OsCYP71D8L-OE* and *cyp71d8l* showed GA deficit phenotypes, we measured the contents of endogenous GAs in *OsCYP71D8L-OE* seedlings, finding that the levels of GA intermediates such as GA1 (bioactive GA), GA53 and GA19 (upstream precursors of GA1), GA9 (precursors of GA4), were significantly reduced in *OsCYP71D8L*-OE seedlings, but the GA24 content was slightly increased in comparison with those in KOS. The levels of GA1, GA9, GA19 and GA53 in *OsCYP71D8L-*OE plants were decreased to about 48%, 27.6%, 11.2% and 9.1% of the WT level, respectively (Fig. 4A), indicating that OsCYP71D8L participated in the deactivation of bioactive GAs in rice.

**Figure 4.**
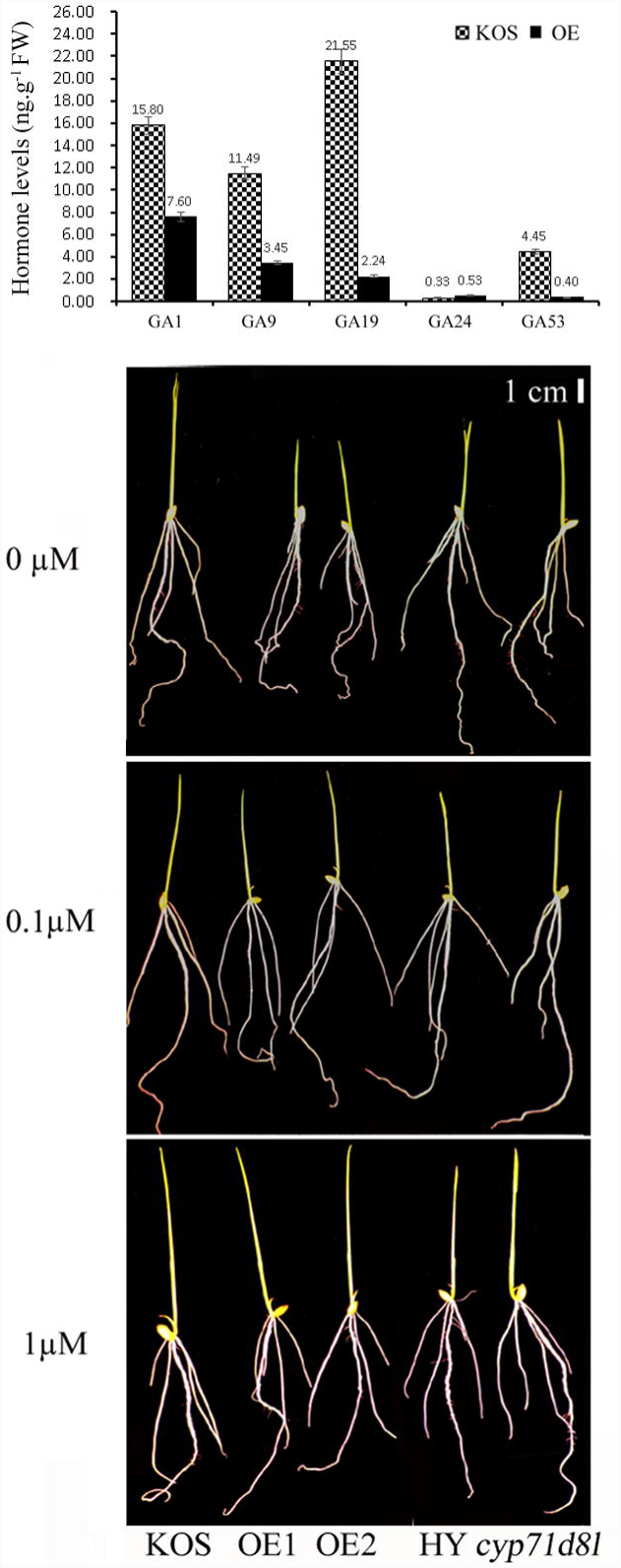
OsCYP71D8L affects GAs homeostasis. A, Measurement of GAs in 4-week-old *OsCYP71D8L-OE* and KOS seedlings. Error bars indicate ± SE (*n*=3). (B) The phenotypes of 6-day-old *OsCYP71D8L-OE*, KOS, *cyp71d8l* and HY seedlings grown in water with 0, 0.1 and 1 µM GA_3_, bar=1cm.

Then, we analyzed the growth of *OsCYP71D8L-OE* and *cyp71d8l* in response to GA3 treatment. Under normal condition, *OsCYP71D8L-OE* seedlings was shorter than KOS, and *OsCYP71D8L-OE* and *cyp71d8l* seedlings displayed shorter root system than that of WT. When treated with 0.1 and 1 µM GA3 for 6 days, the plant height of *OsCYP71D8L-OE* was rescued to that of KOS. Similarly, their root system was also restored to that of KOS and HY with GA3 application, respectively (Fig. 4B). These results strongly suggest that OsCYP71D8L negatively affects GAs level and leads to the growth retardation in *OsCYP71D8L-OE* and *cyp71d8l*.

### OsCYP71D8L affects cytokinin homeostasis and root sensitivity to 6-BA

CK and GA antagonistically control meristem maintenance and organ development (De Smet et al., 2015; Pavlu et al., 2018; Shani et al., 2006), and their levels are counter-balanced by the feedback regulation of GA biosynthesis in rice (Wu et al., 2016). *OsCKX4*, encoding a cytokinin oxidase, is responsible for cytokinin degradation and plays a positive role in crown root formation (Gao et al., 2014). We found that its expression was downregulated in *OsCYP71D8L*-OE and *cyp71d8l* seedlings (Fig 5A), perhaps leading to cytokinin accumulation. Likewise, analysis of endogenous CKs levels including isopentenyladenine (IP), cis-zeatin (CZ) and dihydrozeatin (DZ) showed that the IP content in *OsCYP71D8L-*OE seedlings was increased two folds of KOS (Fig 5B), indicating that cytokinin activity was elevated in *OsCYP71D8L*-OE.

**Figure 5.**
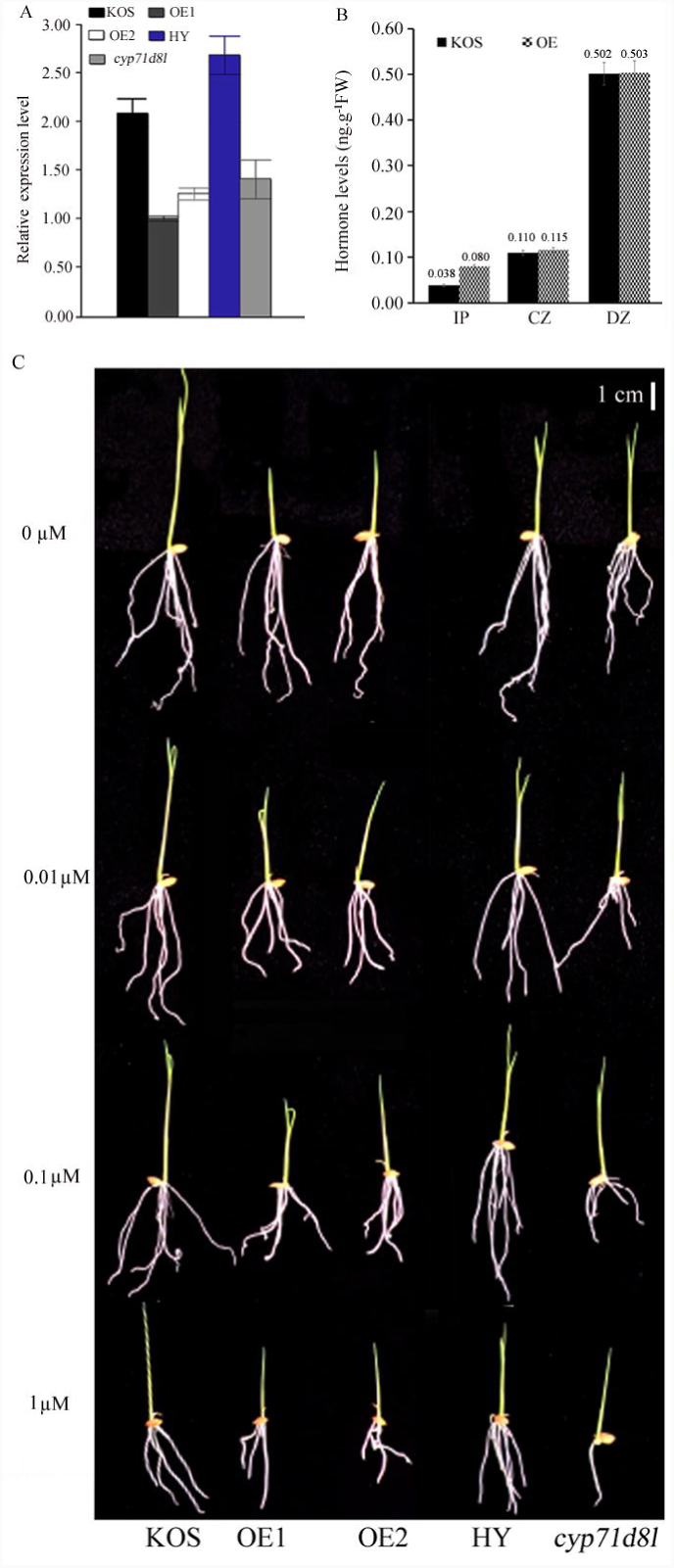
OsCYP71D8L affects CKs homeostasis. A, Relative expression levels of *OsCKX4* in *OsCYP71D8L-OE* and *cyp71d8l* seedlings. B, Measurement of CKs in 4-week-old *OsCYP71D8L-OE* and KOS seedlings. Error bars indicate ± SE (*n*=3).C, The phenotypes of 6-day-old *OsCYP71D8L-OE*, KOS, *cyp71d8l* and HY seedlings treated with 0, 0.001, 0.1 and 1 µM 6-BA, bar=1cm.

We then checked the effect of exogenous 6-Benzylaminopurine (6-BA, a synthetic CK) on the root growth in *OsCYP71D8L*-OE and *cyp71d8l* seedlings. Unlike the root growth promoted by GA, the less developed root system of *OsCYP71D8L*-OE and *cyp71d8l* was more exacerbated by 6-BA than WT. Their root elongation and crown root initiation were notably suppressed with the treatment of 0.001, 0.1 and 1 µM 6-BA (Fig 5C), indicating that the CK over-accumulation is responsible for the under-developed root systems in *OsCYP71D8L*-OE and *cyp71d8l*. Taken together, we proposed that the abnormal GAs and CKs changes caused the dwarf phenotype and less developed root system in *OsCYP71D8L*-OE and *cyp71d8l*.

### OsCYP71D8L positively regulates rice tolerance to drought and salinity stress

Rice growth and development was affected by adverse environment. Overexpression of *PtCYP714A3, OsDSS1* and *CYP94C2b* could improve rice tolerance to drought and salt stress (Kurotani et al., 2015; Tamiru et al., 2015; Wang et al., 2016). Here, we confirmed that *OsCYP71D8L* conferred rice with better tolerance to drought and salinity stress. When 3-week-old seedlings were exposed to drought for 8 days, WT plants exhibited severe stress symptoms with withering leaves, while *OsCYP71D8L-OE* lines and *cyp71d8l* remained normal turgor, and showed mild stress symptoms at the 10^th^ day. After re-watering for 7 days, *OsCYP71D8L-OE* lines and *cyp71d8l* recovered better growth status, while almost all WT plants were dead, (Fig.6A). Similarly, when 3-week-old seedlings treated with 1% NaCl for 7 days, KOS and HY exhibited more severe tissue damages with withering or dead leaves than *OsCYP71D8L*-OE and *cyp71d8l*. After recovery for 7 days, *OsCYP71D8L-OE* and *cyp71d8l* plants recovered fresh green leaves, while WT plants were dead or still withered (Fig.6B).

**Figure 6.**
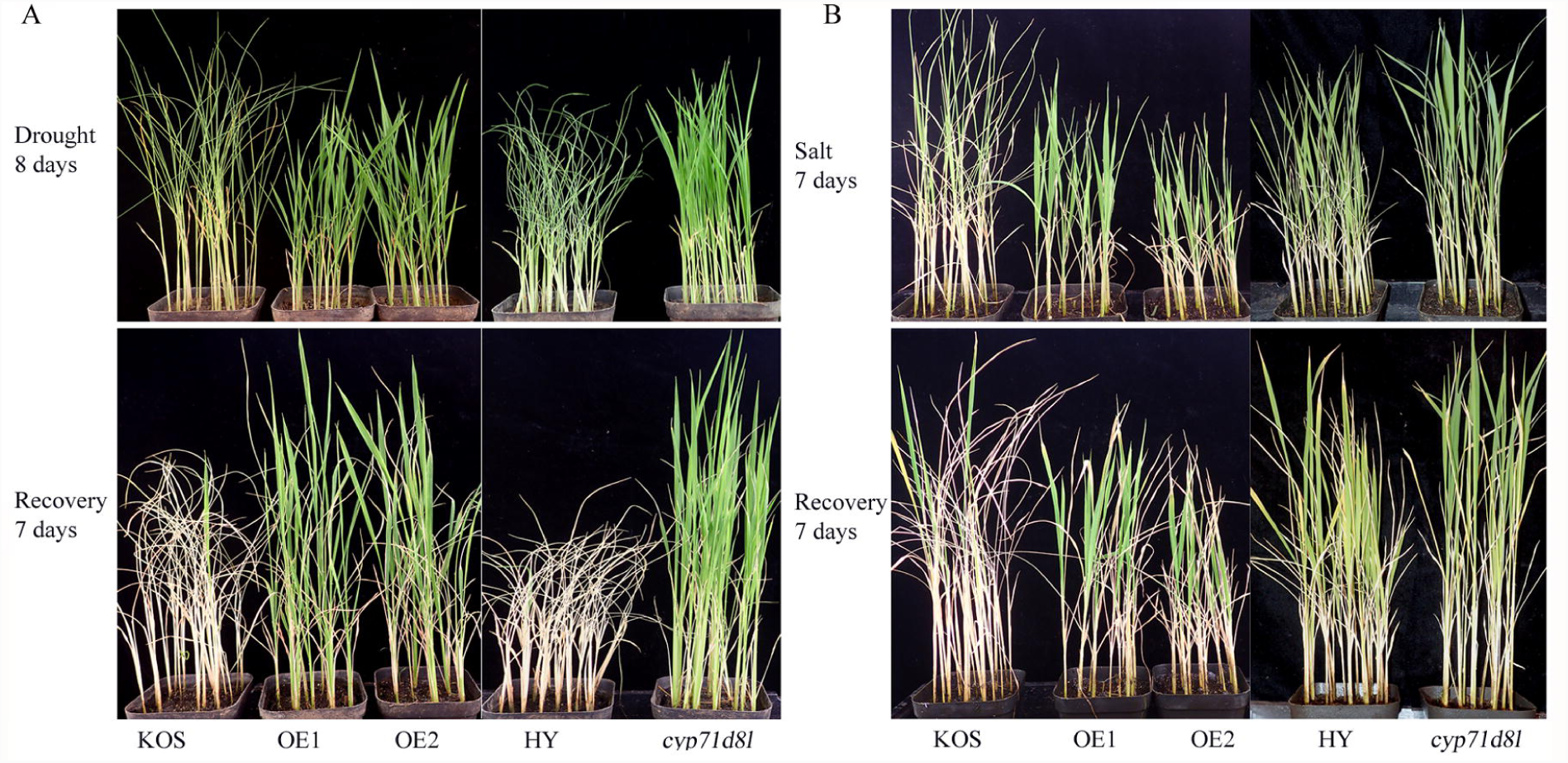
Increased tolerance to drought and salt stress in *OsCYP71D8L*-OE and *cyp71d8l* seedlings. 3-week-old seedlings were treated with drought 10 days or 150 mM NaCl for 7days and followed by re-watering 7 days. Pictures were taken at the 8^th^ day for drought (A), 7^th^ day for salt stress (B) and after recovery 7 days.

### OsCYP71D8L affects chlorophyll and ROS accumulation under stress

To elucidate the physiological characteristics underlaying enhanced tolerance to drought and salt stress, we investigated the contents of soluble sugar and chlorophyll in 3-week-old seedlings. Under normal condition, the levels of soluble sugar in *OsCYP71D8L*-OEs and *cyp71d8l* were about 10% higher than that in KOS and HY, respectively (Fig.7A). Similarly, the levels of chlorophyll a, chlorophyll b and total chlorophyll in *OsCYP71D8L*-OEs and *cyp71d8l* were higher than that in WT under normal condition, respectively. Under drought and salt stress, the levels of total chlorophyll, chlorophyll a and chlorophyll b were decreased in all resulted materials, but remained at higher levels in *OsCYP71D8L*-OE lines and *cyp71d8l* than that in WT (Fig.7B). These results suggested that higher accumulation of soluble sugars and chlorophylls in *OsCYP71D8L*-OE lines and *cyp71d8* might contribute to their enhanced tolerance to drought and salt stress.

**Figure 7.**
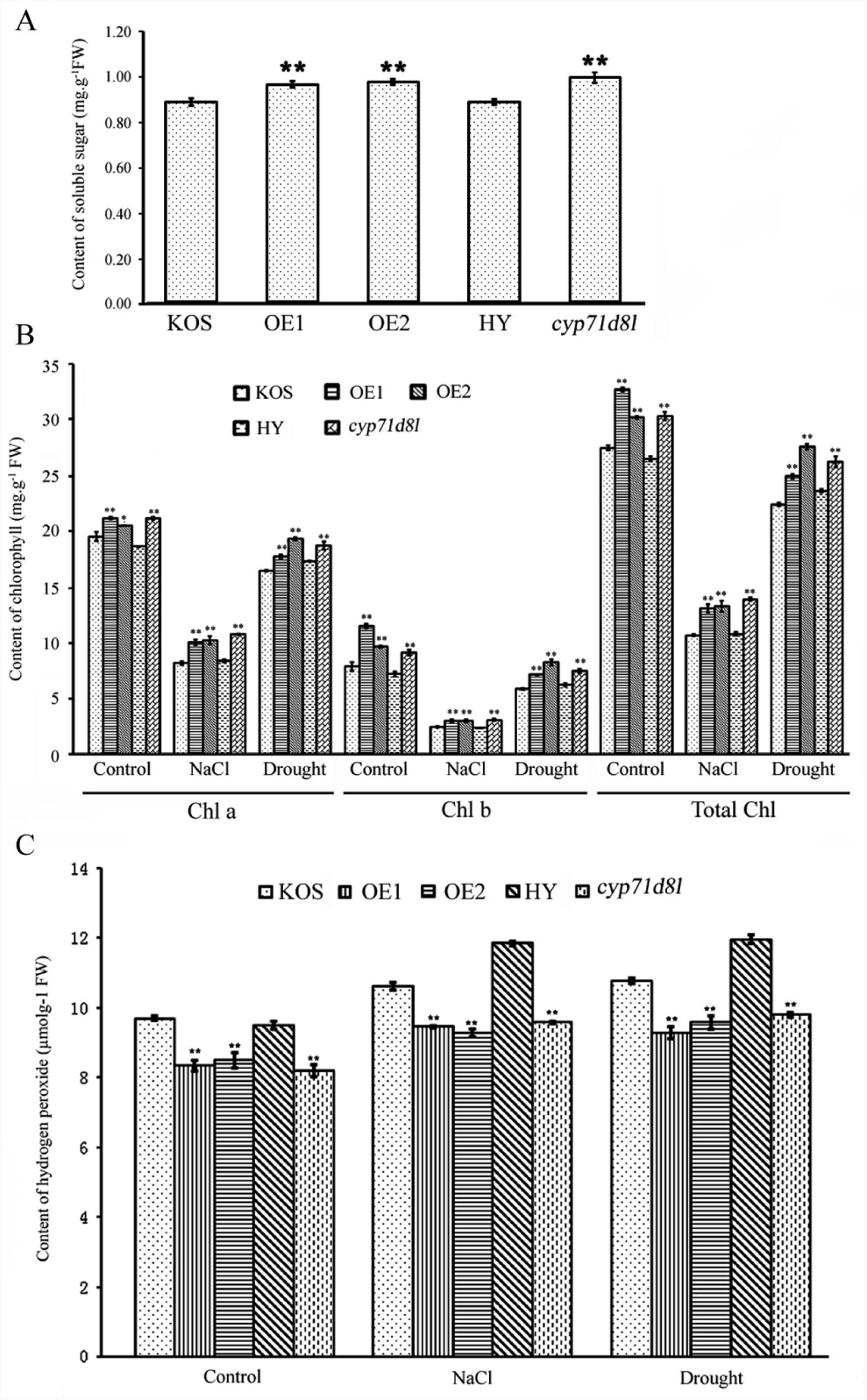
Soluble sugar, chlorophyll, and H_2_O_2_ contents in *OsCYP71D8L*-OE, KOS, *cyp71d8l* and HY seedlings under drought and salt stress. A, Soluble sugar contents in the leaves of 3-week-old seedlings under normal growth condition; B-C, chlorophyll a, chlorophyll b and total chlorophyll (B) and H_2_O_2_ (C) contents upon drought and salt stress. 3-week-old seedlings were subjected to drought for 5 days or 1% NaCl treatment for 3 days.

Exclusive ROS was harmful components seriously affecting plant growth and response to various stresses (del Rio, 2015). In this study, we found that the content of H_2_O_2_ in *OsCYP71D8L*-OE lines and *cyp71d8l* was low compared with that of WT under normal condition. Under drought and salt stress, the H_2_O_2_ level in *OsCYP71D8L*-OE lines and *cyp71d8l* was also significantly lower than that in WT (Fig. 7C), which were coincided with their enhanced tolerance to drought and high salinity.

## Discussion

Plant height is an important trait determining rice architecture and yield, and controlled by a complex phytohormone signaling network. Based on the genetic analysis of dwarf mutants and corresponding genes, GA homeostasis and signaling paly crucial roles in the control of plant height and responses to various stresses in rice (Liu et al., 2018; Wang and Li, 2008). CYP450s catalyze most of the rate-limiting reactions involved in the biosynthesis and catabolism of phytohormones, especially in GAs homeostasis (Hedden and Thomas, 2012; Renault et al., 2014). Rice genome contains at least 356 P450 genes grouped into 10 different clans, and CYP71 clan is the largest set (Nelson and Werck-Reichhart, 2011; Nelson et al., 2004). In sunflowers and sorghum bicolor, several CYP71 members participated in the synthesis of eupatolide, inunolide and sorgoleone (Frey et al., 2018; Pan et al., 2018). In rice, eight CYP450 genes including three highly homologous proteins of CYP71 clan (*OsCYP71D8L, OsCYP71D8* and *OsCYP71D7*) are clustered as a tandem array on chromosome 2, but their characters and functions have been little known. Here, we reported that OsCYP71D8L functioned as a key regulator involved in the regulation of rice growth and development as well as abiotic stress response by affecting the homeostasis of GA and CK. Moreover, the transgenic rice overexpressing *OsCYP71D8L* and its C-terminal truncated mutant *cyp71d8l* exhibited similar phenotypes including dwarf plant, short panicle, reduced grain number per panicle, and enhanced tolerance to drought and salt stress compared with WT, suggesting that *cyp71d8l* was gain-of-function mutant.

In rice, multiple CYP40s not belonging to the CYP71 clan were reported to regulate plant developmental processes and stress response by modulating GA homoeostasis (Itoh et al., 2004; Magome et al., 2013; Tamiru et al., 2015; Wang et al., 2012; Zhang et al., 2008; Zhu et al., 2006). Here, we confirmed that overexpression of *OsCYP71D8L* caused the notably decreased accumulation of GA1, GA9, GA19 and GA53. Consistent with this, *OsCYP71D8L*-OE and *cyp71d8l* exhibited the dwarf phenotypes with reduced node and internode number, and exogenous GA3 could rescue their plant height, suggesting that OsCYP71D8L is involved in the control of plant height by affecting GA homeostasis in rice.

CKs and GAs antagonistically regulate plant shoot and root growth by mediating meristem activity and cell division (Azizi et al., 2015; Kyozuka, 2007; Pacifici et al., 2015), and their homeostasis are counter-balanced by the feedback regulation of GA biosynthesis in rice (Wu et al., 2016). Generally, GAs has positive effect on shoot and root elongation in rice. For examples, AP2/ERF transcription factor SHB controls root elongation through dose-dependent effects of GA on cell elongation and proliferation (Li et al.2015). GDD1 mediates cell elongation by regulating GA biosynthesis, and *gdd1* showed gibberellin-deficient phenotypes with reduced length of root, stems, spikes, and seeds (Li et al. 2011). Similarly, *OsMCA1* is involved in the deactivation of bioactive GA, and its mutation results in GA-deficient phenotypes with shorter root and reduced height (Liu et al. 2015). In rice, cytokinin is essential for the control of entire life cycle including root development, shoot meristem activity, vegetative and reproductive branching, spikelets per panicle, and grain production(Ashikari et al., 2005; Du et al., 2017; Gao et al., 2014; Gu et al., 2015; Jiang et al., 2017; Kurakawa et al., 2007; Li et al., 2013; Ohashi et al., 2017; Panda et al., 2018; Wu et al., 2017; Wu et al., 2016; Zhao et al., 2015). For examples, cytokinin receptor OsCKT1 is necessary for the regulation of root development, and *Osckt1* exhibits insensitivity to cytokinin treatment with normal root growth (Ding et al., 2017). ERF3 and WOX11 regulates crown root development by tuning auxin- and cytokinin-signaling (Zhao et al., 2015). Overexpression of *KNOTTED1-like* results in high-CK and low-GA and inhibits root development by inducting *OsIPT2* and *OsIPT3* expression (Sakamoto et al., 2006). Similarly, down-regulation of CK signaling leads to reduced internode lengths and tiller number, and promoted lateral root growth (Sun et al., 2014). Thus, the appropriate GAs and CKs homeostasis is required for the normal growth and development in rice. In this study, we confirmed that overexpression of *OsCYP71D8L* led to the low-GA and high-CK associated with the abnormal root system and dwarf plants. Moreover, the less developed root system in *OsCYP71D8L*-OE and *cyp71d8l* could be rescued by GA3, but exacerbated by 6-BA. At the transcription level, *OsCKX4* expression was down-regulated in *OsCYP71D8L-OE* and *cyp71d8l* seedlings, which catalyzed the degradation of cytokinins and mediated crown root development and plant height by integrating CK and auxin signaling (Gao et al., 2014). Its down-regulation might partially account for the elevated CK activity and the abnormal phenotypes in *OsCYP71D8L*-OE and *cyp71d8l*.

GA and CK are crucial in promoting cell division, cell growth and differentiation, and have large impact on yield-related traits in crops (Kyozuka, 2007; Nadolska-Orczyk et al., 2017). Two green revolution genes, wheat Reduced height1 (*Rht1*) and rice semi-dwarf1 (*sd1*), are involved in GA signaling and biosynthesis, suggesting that manipulating GA level or signaling is a most effective approach for producing high-yield cultivars (Sakamoto, 2006). In rice, moderately reduced GA levels by ectopic expressing *GA2ox6* mutants caused moderate plant height, more tillers, and high grain yield, but excessively reduced GA levels resulted in over-dwarfing plants and serious yield decrease (Lo et al., 2017). In comparison, it is also feasible to genetically improve crop yield by modulating CK homeostasis. For examples, exogenous CK could increase rice yield potential by enhancing grain filling and bearing large panicles (Panda et al., 2018). Downregulation of *OsCKX2* and *OscZOG1* resulted in elevated CKs, and thereby improved yield-related traits including tiller numbers, panicle branches, and grain number per panicle in rice (Ashikari et al., 2005; Shang et al., 2016). Similarly, GA20ox1 affected grain number and yield by rebalancing activities of CKs and GAs in rice (Wu et al., 2016). However, An-2, as a cytokinin synthesis enzyme, could increase endogenous cytokinin content, but decreased grain number per panicle and tiller number (Gu et al., 2015). Thus, the suitable GAs and CKs homeostasis is essential for the regulation of various yield-related traits in rice. Our results confirmed that overexpression of *OsCYP71D8L* resulted in the significantly lowered GAs and moderately elevated CKs levels, which thereby led to the reduced panicle length and grain number per panicle in *OsCYP71D8L*-OE and *cyp71d8l* plants.

Phytohormones and their crosstalk play central roles in coordinating plant response to adverse environments. Recent data has shown that the reduction of GA synthesis and signaling is conducive to plant adaptation to abiotic stresses including drought, cold, salt and osmotic stress (Colebrook et al., 2014). For examples, overexpressing *AtGAMT1* reduced the active GAs levels, but enhanced drought tolerance in tomato (Nir et al., 2014). In rice, *gid1* (a mutant of a GA receptor) exhibited enhanced submergence tolerance associated with less ROS accumulation and increased chlorophyll and carbohydrates (Du et al., 2015). Ectopic expression of *GA2ox6* mutants and *PtCYP714A3* led to reduced GA levels and dwarfed plant, but conferred rice with enhanced tolerance to drought and salt stress (Lo et al., 2017; Wang et al., 2016). Unlike GAs, CKs positively regulates plant stress tolerance by enhancing photosynthesis and antioxidant capacities, improving water use efficiency, and modulating growth and development (Pavlu et al., 2018). In rice, CKs could delay dark-induced senescence by maintaining the chlorophyll cycle and photosynthetic stability (Talla et al., 2016). Overexpressing *ISOPENTENYLTRANSFERASE* could lead to high CKs levels and improve drought tolerance by coordinating source/sink relationships and nutrient metabolism in rice (Peleg et al., 2011; Reguera et al., 2013). Similarly, OsAHP1 and OsAHP2 function as positive regulators of the cytokinin signaling and are required for the regulation salt and drought tolerance in rice (Sun et al., 2014). These findings show that GA and CK play antagonistic roles in plant response to abiotic stresses, and manipulating GA and CK levels is an effective strategy to improve stress tolerance in rice. Coincided with previous studies, overexpression of *OsCYP71D8L* caused low-GA and high-CK levels. These hormone changes might confer *OsCYP71D8L*-OE and *cyp71d8l* with improved tolerance to drought and salt stress by maintaining high levels of chlorophyll and soluble sugar, and low ROS accumulation.

The growth readjustment and adaptation to various stresses are key to the survival of plants under adverse environments. Our study confirmed that OsCYP71D8L, as a potential GA-deactivating protein, might act as key regulator balancing the growth process and stress response by coordinating GA and CK homeostasis. These findings also provide important information for engineering stress-tolerant rice by modifying *OsCYP71D8L* with genome editing technology or ectopic expressing *OsCYP71D8L* driven by stress inducible promoters.

## Acknowledgements

This work was supported by grants from the National Key Research and Development Program of China (2016YFD0100901) and National Natural Science Foundation of China (31701781).

---

**Figure S1 Amino acid sequences homologous analysis of OsCYP71D8L, OsCYp71D8, LOC4328533, LOC4328535 and OsCYP71D7.** Alignments were performed using DNAMAN (version 8.0).

**Figure S2. Position and sequence analyses of the T-DNA insertion in *cyp71d8l*.** A, Position of the T-DNA insertion in the *OsCYP71D8L* and genotype test. Genotype of *cyp71d8l* was showed in the C-terminal of *OsCYP71D8L* based on genomic DNA. PCR test was performed using 3 primers (Left primer 5-AGACAAGTGGACATCCGCTC-3, Right primer 5-AGGTTACATCCTGTGGTGCC-3, and RB primer located in the pGA2707 vector 5-GTTACG TCCTGTAGAAACCCCAA-3). B-C, The T-DNA insertion resulted in the deletion of 93 bases and the truncated protein of 30 amino acids at C-terminal of *OsCYP71d8L*. DNA sequence alignment analysis of PCR product from (A), OsCYP71D8L and pGA2772 vector.

**Figure S3. Phenotype of *OsCYP71D8*L-OE and *cyp71d8l*. A,** Seven-day-old seedlings of *OsCYP71D8*L-OE, *cyp71d8l*, KOS and HY. B, Seven-week-old plants of *OsCYP71D8*L-OE, *cyp71d8l*, KOS and HY. C, Plants of *OsCYP71D8*L-OE, *cyp71d8l*, KOS and HY at the flowering stage.

## References

Ashikari, M., Sakakibara, H., Lin, S.Y., Yamamoto, T., Takashi, T., Nishimura, A., Angeles, E.R., Qian, Q., Kitano, H. and Matsuoka, M. (2005) Cytokinin oxidase regulates rice grain production. Science 309, 741–745.

Azizi, P., Rafii, M.Y., Maziah, M., Abdullah, S.N., Hanafi, M.M., Latif, M.A., Rashid, A.A. and Sahebi, M. (2015) Understanding the shoot apical meristem regulation: a study of the phytohormones, auxin and cytokinin, in rice. Mech Develop. 135, 1–15.

Colebrook, E.H., Thomas, S.G., Phillips, A.L. and Hedden, P. (2014) The role of gibberellin signalling in plant responses to abiotic stress. J Exp Biol 217, 67–75.

De Smet, S., Cuypers, A., Vangronsveld, J. and Remans, T. (2015) gene networks involved in hormonal control of root development in Arabidopsis thaliana: a framework for studying its disturbance by metal stress. Int. J. Mol. Sci. 16, 19195–19224.

del Rio, L.A. (2015) ROS and RNS in Plant physiol.: an overview. J. Exp. Bot. 66, 2827–2837.

Ding, W.N., Tong, H.S., Zheng, W.J., Ye, J., Pan, Z.C., Zhang, B.T. and Zhu, S.H. (2017) Isolation, characterization and transcriptome analysis of a cytokinin receptor mutant Osckt1 in rice. Front. Plant Sci. 8.

Du, H., Chang, Y., Huang, F. and Xiong, L.Z. (2015) GID1 modulates stomatal response and submergence tolerance involving abscisic acid and gibberellic acid signaling in rice. J Integr. Plant Biol. 57, 954–968.

Du, Y.F., Liu, L., Li, M.F., Fang, S., Shen, X.M., Chu, J.F. and Zhang, Z.X. (2017) UNBRANCHED3 regulates branching by modulating cytokinin biosynthesis and signaling in maize and rice. New Phytologist 214, 721–733.

Frey, M., Schmauder, K., Pateraki, I. and Spring, O. (2018) Biosynthesis of eupatolide-a metabolic route for sesquiterpene lactone formation involving the P450 enzyme CYP71DD6. Acs Chem Biol 13, 1536–1543.

Gao, S., Fang, J., Xu, F., Wang, W., Sun, X., Chu, J., Cai, B., Feng, Y. and Chu, C. (2014) CYTOKININ OXIDASE/DEHYDROGENASE4 integrates cytokinin and auxin signaling to control rice crown root formation. Plant Physiol. 165, 1035–1046.

Gu, B.G., Zhou, T.Y., Luo, J.H., Liu, H., Wang, Y.C., Shangguan, Y.Y., Zhu, J.J., Li, Y., Sang, T., Wang, Z.X. and Han, B. (2015) an-2 encodes a cytokinin synthesis enzyme that regulates awn length and grain production in rice. Mol. plant 8, 1635–1650.

Itoh, H., Tatsumi, T., Sakamoto, T., Otomo, K., Toyomasu, T., Kitano, H., Ashikari, M., Ichihara, S. and Matsuoka, M. (2004) A rice semi-dwarf gene, Tan-Ginbozu (D35), encodes the gibberellin biosynthesis enzyme, ent-kaurene oxidase. Plant Mol. Biol. 54, 533–547.

Jiang, W., Zhou, S.L., Zhang, Q., Song, H.Z., Zhou, D.X. and Zhao, Y. (2017) Transcriptional regulatory network of WOX11 is involved in the control of crown root development, cytokinin signals, and redox in rice. J. Exp. Bot. 68, 2787–2798.

Kurakawa, T., Ueda, N., Maekawa, M., Kobayashi, K., Kojima, M., Nagato, Y., Sakakibara, H. and Kyozuka, J. (2007) Direct control of shoot meristem activity by a cytokinin-activating enzyme. Nature 445, 652–655.

Kurotani, K., Hayashi, K., Hatanaka, S., Toda, Y., Ogawa, D., Ichikawa, H., Ishimaru, Y., Tashita, R., Suzuki, T., Ueda, M., Hattori, T. and Takeda, S. (2015) Elevated levels of CYP94 family gene expression alleviate the jasmonate response and enhance salt tolerance in rice. Plant Cell Physiol. 56, 779–789.

Kwon, C.T. and Paek, N.C. (2016) Gibberellic Acid: A Key Phytohormone for Spikelet Fertility in Rice Grain Production. Int. J. Mol. Sci. 17.

Kyozuka, J. (2007) Control of shoot and root meristem function by cytokinin. Curr Opin Plant Biol 10, 442–446.

Li, J.A., Jiang, J.F., Qian, Q.A., Xu, Y.Y., Zhang, C., Xiao, J., Du, C., Luo, W., Zou, G.X., Chen, M.L., Huang, Y.Q., Feng, Y.Q., Cheng, Z.K., Yuan, M. and Chong, K. (2011) Mutation of Rice BC12/GDD1, which encodes a kinesin-like protein that binds to a ga biosynthesis gene promoter, leads to dwarfism with impaired cell elongation. Plant Cell 23, 628–640.

Li, S.Y., Zhao, B.R., Yuan, D.Y., Duan, M.J., Qian, Q., Tang, L., Wang, B., Liu, X.Q., Zhang, J., Wang, J., Sun, J.Q., Liu, Z., Feng, Y.Q., Yuan, L.P. and Li, C.Y. (2013) Rice zinc finger protein DST enhances grain production through controlling Gn1a/OsCKX2 expression. Proc. Natl Acad. Sci. USA 110, 3167–3172.

Liu, F., Wang, P.D., Zhang, X.B., Li, X.F., Yan, X.H., Fu, D.H. and Wu, G. (2018) The genetic and molecular basis of crop height based on a rice model. Planta 247, 1–26.

Liu, Z.W., Cheng, Q., Sun, Y.F., Dai, H.X., Song, G.Y., Guo, Z.B., Qu, X.F., Jiang, D.M., Liu, C., Wang, W. and Yang, D.C. (2015) A SNP in OsMCA1 responding for a plant architecture defect by deactivation of bioactive GA in rice. Plant Mol. Biol. 87, 17–30.

Lo, S.F., Ho, T.H.D., Liu, Y.L., Jiang, M.J., Hsieh, K.T., Chen, K.T., Yu, L.C., Lee, M.H., Chen, C.Y., Huang, T.P., Kojima, M., Sakakibara, H., Chen, L.J. and Yu, S.M. (2017) Ectopic expression of specific GA2 oxidase mutants promotes yield and stress tolerance in rice. Plant Biotech. J. 15, 850–864.

Magome, H., Nomura, T., Hanada, A., Takeda-Kamiya, N., Ohnishi, T., Shinma, Y., Katsumata, T., Kawaide, H., Kamiya, Y. and Yamaguchi, S. (2013) CYP714B1 and CYP714B2 encode gibberellin 13-oxidases that reduce gibberellin activity in rice. Proc. Natl Acad. Sci. USA 110, 1947–1952.

Nadolska-Orczyk, A., Rajchel, I.K., Orczyk, W. and Gasparis, S. (2017) Major genes determining yield-related traits in wheat and barley. Theor Appl Genet 130, 1081–1098.

Nelson, D. and Werck-Reichhart, D. (2011) A P450-centric view of plant evolution. Plant J. 66, 194–211.

Nelson, D.R., Schuler, M.A., Paquette, S.M., Werck-Reichhart, D. and Bak, S. (2004) Comparative genomics of rice and Arabidopsis analysis of 727 cytochrome P450 genes and pseudogenes from a monocot and a dicot. Plant Physiol. 135, 756–772.

Nir, I., Moshelion, M. and Weiss, D. (2014) The Arabidopsis GIBBERELLIN METHYL TRANSFERASE 1 suppresses gibberellin activity, reduces whole-plant transpiration and promotes drought tolerance in transgenic tomato. Plant Cell &Environ. 37, 113–123.

Nomura, T., Magome, H., Hanada, A., Takeda-Kamiya, N., Mander, L.N., Kamiya, Y. and Yamaguchi, S. (2013) Functional analysis of Arabidopsis CYP714A1 and CYP714A2 reveals that they are distinct gibberellin modification enzymes. Plant Cell Physiol. 54, 1837–1851.

Ohashi, M., Ishiyama, K., Kojima, S., Kojima, M., Sakakibara, H., Yamaya, T. and Hayakawa, T. (2017) Lack of cytosolic glutamine synthetase1;2 activity reduces nitrogen-dependent biosynthesis of cytokinin required for axillary bud outgrowth in rice seedlings. Plant Cell Physiol. 58, 679–690.

Pacifici, E., Polverari, L. and Sabatini, S. (2015) Plant hormone cross-talk: the pivot of root growth. J. Exp. Bot. 66, 1113–1121.

Pan, Z.Q., Baerson, S.R., Wang, M., Bajsa-Hirschel, J., Rimando, A.M., Wang, X.Q., Nanayakkara, N.P.D., Noonan, B.P., Fromm, M.E., Dayan, F.E., Khan, I.A. and Duke, S.O. (2018) A cytochrome P450 CYP71 enzyme expressed in Sorghum bicolor root hair cells participates in the biosynthesis of the benzoquinone allelochemical sorgoleone. New Phytologist 218, 616–629.

Panda, B.B., Sekhar, S., Dash, S.K., Behera, L. and Shaw, B.P. (2018) Biochemical and molecular characterisation of exogenous cytokinin application on grain filling in rice. BMC Plant Biol. 18.

Pavlu, J., Novak, J., Koukalova, V., Luklova, M., Brzobohaty, B. and Cerny, M. (2018) cytokinin at the crossroads of abiotic stress signalling pathways. Int. J. Mol. Sci. 19.

Peleg, Z., Reguera, M., Tumimbang, E., Walia, H. and Blumwald, E. (2011) Cytokinin-mediated source/sink modifications improve drought tolerance and increase grain yield in rice under water-stress. Plant Biotech. J. 9, 747–758.

Regnault, T., Daviere, J.M., Heintz, D., Lange, T. and Achard, P. (2014) The gibberellin biosynthetic genes AtKAO1 and AtKAO2 have overlapping roles throughout Arabidopsis development. Plant J. 80, 462–474.

Reguera, M., Peleg, Z., Abdel-Tawab, Y.M., Tumimbang, E.B., Delatorre, C.A. and Blumwald, E. (2013) Stress-induced cytokinin synthesis increases drought tolerance through the coordinated regulation of carbon and nitrogen assimilation in rice. Plant Physiol. 163, 1609–1622.

Renault, H., Bassard, J.E., Hamberger, B. and Werck-Reichhart, D. (2014) Cytochrome P450-mediated metabolic engineering: current progress and future challenges. Curr Opin Plant Biol 19, 27–34.

Sakamoto, T. (2006) Phytohormones and rice crop yield: strategies and opportunities for genetic improvement. Transgenic Res. 15, 399–404.

Sakamoto, T., Sakakibara, H., Kojima, M., Yamamoto, Y., Nagasaki, H., Inukai, Y., Sato, Y. and Matsuoka, M. (2006) Ectopic expression of KNOTTED1-like homeobox protein induces expression of cytokinin biosynthesis genes in rice. Plant Physiol. 142, 54–62.

Shang, X.L., Xie, R.R., Tian, H., Wang, Q.L. and Guo, F.Q. (2016) Putative zeatin O-glucosyltransferase OscZOG1 regulates root and shoot development and formation of agronomic traits in rice. J. Integr. Plant Biol. 58, 627–641.

Shani, E., Yanai, O. and Ori, N. (2006) The role of hormones in shoot apical meristem function. Curr Opin Plant Biol 9, 484–489.

Song, S.Y., Chen, Y., Chen, J., Dai, X.Y. and Zhang, W.H. (2011) Physiological mechanisms underlying OsNAC5-dependent tolerance of rice plants to abiotic stress. Planta 234, 331–345.

Sun, L.J., Zhang, Q., Wu, J.X., Zhang, L.Q., Jiao, X.W., Zhang, S.W., Zhang, Z.G., Sun, D.Y., Lu, T.G. and Sun, Y. (2014) Two rice authentic histidine phosphotransfer proteins, OsAHP1 and OsAHP2, mediate cytokinin signaling and stress responses in rice. Plant Physiol. 165, 335–345.

Talla, S.K., Panigrahy, M., Kappara, S., Nirosha, P., Neelamraju, S. and Ramanan, R. (2016) Cytokinin delays dark-induced senescence in rice by maintaining the chlorophyll cycle and photosynthetic complexes. J. Exp. Bot. 67, 1839–1851.

Tamiru, M., Undan, J.R., Takagi, H., Abe, A., Yoshida, K., Undan, J.Q., Natsume, S., Uemura, A., Saitoh, H., Matsumura, H., Urasaki, N., Yokota, T. and Terauchi, R. (2015) A cytochrome P450, OsDSS1, is involved in growth and drought stress responses in rice (Oryza sativa L.). Plant Mol. Biol. 88, 85–99.

Tanaka, N., Matsuoka, M., Kitano, H., Asano, T., Kaku, H. and Komatsu, S. (2006) gid1, a gibberellin-insensitive dwarf mutant, shows altered regulation of probenazole-inducible protein (PBZ1) in response to cold stress and pathogen attack. Plant Cell & Environ. 29, 619–631.

Ueguchi-Tanaka, M., Ashikari, M., Nakajima, M., Itoh, H., Katoh, E., Kobayashi, M., Chow, T.Y., Hsing, Y.I.C., Kitano, H., Yamaguchi, I. and Matsuoka, M. (2005) GIBBERELLIN INSENSITIVE DWARF1 encodes a soluble receptor for gibberellin. Nature 437, 693–698.

Urano, D., Colaneri, A. and Jones, A.M. (2014) G alpha modulates salt-induced cellular senescence and cell division in rice and maize. J. Exp. Bot. 65, 6553–6561.

Wang, C.T., Yang, Y., Wang, H.H., Ran, X.J., Li, B., Zhang, J.T. and Zhang, H.X. (2016) Ectopic expression of a cytochrome P450 monooxygenase gene PtCYP714A3 from Populus trichocarpa reduces shoot growth and improves tolerance to salt stress in transgenic rice. Plant Biotech. J. 14, 1838–1851.

Wang, Q., Hillwig, M.L., Wu, Y.S. and Peters, R.J. (2012) CYP701A8: a rice ent-kaurene oxidase paralog diverted to more specialized diterpenoid metabolism. Plant Physiol. 158, 1418–1425.

Wang, Y.H. and Li, J.Y. (2008) Molecular basis of plant architecture. Annu Rev Plant Biol. 59, 253–279.

Wu, C., Cui, K.H., Wang, W.C., Li, Q., Fahad, S., Hu, Q.Q., Huang, J.L., Nie, L.X., Mohapatra, P.K. and Peng, S.B. (2017) Heat-induced cytokinin transportation and degradation are associated with reduced panicle cytokinin expression and fewer spikelets per panicle in rice. Front. Plant Sci. 8.

Wu, Y., Wang, Y., Mi, X.F., Shan, J.X., Li, X.M., Xu, J.L. and Lin, H.X. (2016) The QTL GNP1 encodes ga20ox1, which increases grain number and yield by increasing cytokinin activity in rice panicle meristems. PLoS Genet. 12.

Zhang, H.W., Pan, X.W., Li, Y.C., Wan, L.Y., Li, X.X. and Huang, R.F. (2012) Comparison of differentially expressed genes involved in drought response between two elite rice varieties. Mol. Plant 5, 1403–1405.

Zhang, Y.Y., Zhang, B.C., Yan, D.W., Dong, W.X., Yang, W.B., Li, Q., Zeng, L.J., Wang, J.J., Wang, L.Y., Hicks, L.M. and He, Z.H. (2011) Two Arabidopsis cytochrome P450 monooxygenases, CYP714A1 and CYP714A2, function redundantly in plant development through gibberellin deactivation. Plant J. 67, 342–353.

Zhang, Y.Y., Zhu, Y.Y., Peng, Y., Yan, D., Li, Q., Wang, J., Wang, L.Y. and He, Z.H. (2008) Gibberellin homeostasis and plant height control by EUI and a role for gibberellin in root gravity responses in rice. Cell Res. 18, 412–421.

Zhao, Y., Cheng, S.F., Song, Y.L., Huang, Y.L., Zhou, S.L., Liu, X.Y. and Zhou, D.X. (2015) The interaction between rice ERF3 and WOX11 promotes crown root development by regulating gene expression involved in cytokinin signaling. Plant Cell 27, 2469–2483.

Zhu, Y.Y., Nomura, T., Xu, Y.H., Zhang, Y.Y., Peng, Y., Mao, B.Z., Hanada, A., Zhou, H.C., Wang, R.X., Li, P.J., Zhu, X.D., Mander, L.N., Kamiya, Y., Yamaguchi, S. and He, Z.H. (2006) ELONGATED UPPERMOST INTERNODE encodes a cytochrome P450 monooxygenase that epoxidizes gibberellins in a novel deactivation reaction in rice. Plant Cell 18, 442–456.

